# Microsaccades do not give rise to a conscious feeling of agency for their sensorimotor consequences in visual perception

**DOI:** 10.1101/2025.05.04.652094

**Authors:** Jan-Nikolas Klanke, Sven Ohl, Martin Rolfs

## Abstract

Feeling of agency (FoA)—the experience of controlling one’s actions and their outcomes—has been widely studied for bodily movements. Here, we investigated if microsaccades—small ballistic eye movements—are equally characterized by FoA and if intention mediates this sense of control. We measured FoA via intentional binding, a perceived compression between an action and its effect. In our experiments, we presented a vertically oriented grating, rendered invisible during stable fixation by a rapid temporal phase shift (>60 Hz) that became visible when its retinal motion was slowed down by a microsaccade (active condition). The stimulus was embedded in a clock face and observers reported perceived stimulus timing in each trial. Perceived timing of microsaccade-contingent stimulus perception was compared to the replay of a previous microsaccade’s retinal consequence (replay condition). Trials without a stimulus were included as a control. To examine the role of intention, we tested this paradigm across two experiments in which observers were either instructed to saccade (intended microsaccades) or fixate (unintended microsaccades). In Experiment 2, no instruction was administered such that any microsaccades were considered spontaneous. Microsaccades—either actively generated or replayed— consistently rendered the stimulus highly visible compared to trials without such movements— provided microsaccade direction and peak velocity aligned with the stimulus’s motion. Temporal estimates did not differ between the active and replay conditions for any microsaccade type. This result suggests the absence of temporal binding between eye movements and their sensory consequences, and that intention does not facilitate FoA for small eye movements.

**Significance statement:** Eye movements reflect our decision to closer inspect an aspect of the environment—bodily actions that align our perception with a preceding intention. Here, we investigated if microsaccades—a ballistic, minuscule type of saccade—can be characterized by a feeling of agency: the faint experience of affecting change through intentional actions. In two experiments, we presented an identical stimulus whose visibility was either gaze-contingent (active condition) or independent of eye movements (replay condition). In **Experiment 1**, we directly compared intended and unintended microsaccades and contrasted them with spontaneous microsaccades in **Experiment 2**. We found no difference between the active and replay condition for either eye movement type. Our data, hence, does not support feeling of agency for microsaccades. While it remains an open question if large saccades are characterized by feeling of agency, our finding demonstrates that intention is not sufficient to elicit feeling of agency for minuscule motor acts.

## Introduction

When we execute an action, we typically experience the underlying process as a coherent and continuous stream of events from intention via bodily movement to effects in the world. This minimal (Gallagher, 2000) or ‘phenomenally thin’ (Clark et al., 2013; Haggard, 2017) experience of control over our actions and (through them) their consequences (Moore, 2016) is called *feeling of agency* (FoA; Haggard, 2017; Synofzik et al., 2008). Here, we investigate if observers experience FoA for eye movements and their outcomes (i.e., changes in visual information), or if such visual actions (cf. Schweitzer & Rolfs, 2022) cannot be characterized in terms of FoA. Theoretically, such visual FoA (vFOA) should unfold just like FoA for other bodily actions: The observer has a goal (e.g., explore a new aspect of the visual scene), forms the intention to make an eye movement, executes it, and, consequently, is presented with a change in visual information. Nevertheless, the scarcity of phenomenological descriptions of eye movements as actions that elicit a vFoA, along with their peculiar relation to volition (Hogendoorn, 2016), their direct link to cognition (Kennard et al., 2005), high frequency (Wolfe et al., 2011), and minimal proprioceptive feedback (Skavenski et al., 1972; Steinbach, 1987), raises the question of whether observers experience vFoA, or if eye movements occur without the subtle and transient experience of agency.

To investigate this question, we used a specific type of eye movement as actions: microsaccades. These are small, binocularly coordinated, ballistic eye movements (Martinez-Conde et al., 2004; Rucci & Poletti, 2015) with distinctly correlated movement parameters (i.e, amplitude and peak velocity; Zuber et al., 1965), typically executed spontaneously during stable fixation (Rolfs, 2009). Recent behavioral and neurophysiological studies show that microsaccades are generated within the same oculomotor network as larger saccades (Hafed et al., 2009; Hafed, 2011; Otero-Millan et al., 2008; Rolfs et al., 2008) and exert similar functions, for example visual stabilization (Martinez-Conde et al., 2013), enhancement of visual acuity (especially for fine spatial details; Ko et al., 2010; Poletti, 2023), as well as a reduction of binocular disparity (Otero-Millan et al., 2014). It is these structural and functional similarities to large saccades that indicate that microsaccades dispatch efference copies (Bridgeman, 1995; Findlay, 1974) and, hence, bear the potential to influence FoA (cf. Blakemore et al., 2000; Haggard, 2017).

Examining vFoA for eye movements requires that these actions—here, microsaccades—give rise to distinct perceptual consequences. To comply with this demand, we presented a high temporal-frequency stimulus that was rendered invisible by its high internal velocity (>60 Hz, cf. Castet & Masson, 2000). If, however, a microsaccade with matching kinematics occurs, it would briefly stabilize (or, reduce the speed of) the stimulus on the retina (cf. Deubel & Elsner, 1986; Kelly, 1990; **Fig. 1b/c**), and render it visible as a ‘robust, short flash of the seemingly stationary grating’ (Deubel et al., 1987). Thus, the stimulus appeared as an immediate sensory consequence of an activly generated microsaccade (*active condition*). To create the same perceptual impression in the absence of an action, we replayed the visual consequences of a previous eye movement back to the observer: By shifting its aperture, the stimulus was stabilized on the screen similarly to how an active microsaccade stabilized the grating on the retina—but in the absence of an eye movement (*replay condition*; **Fig. 1c/d**).

**Fig 1.**
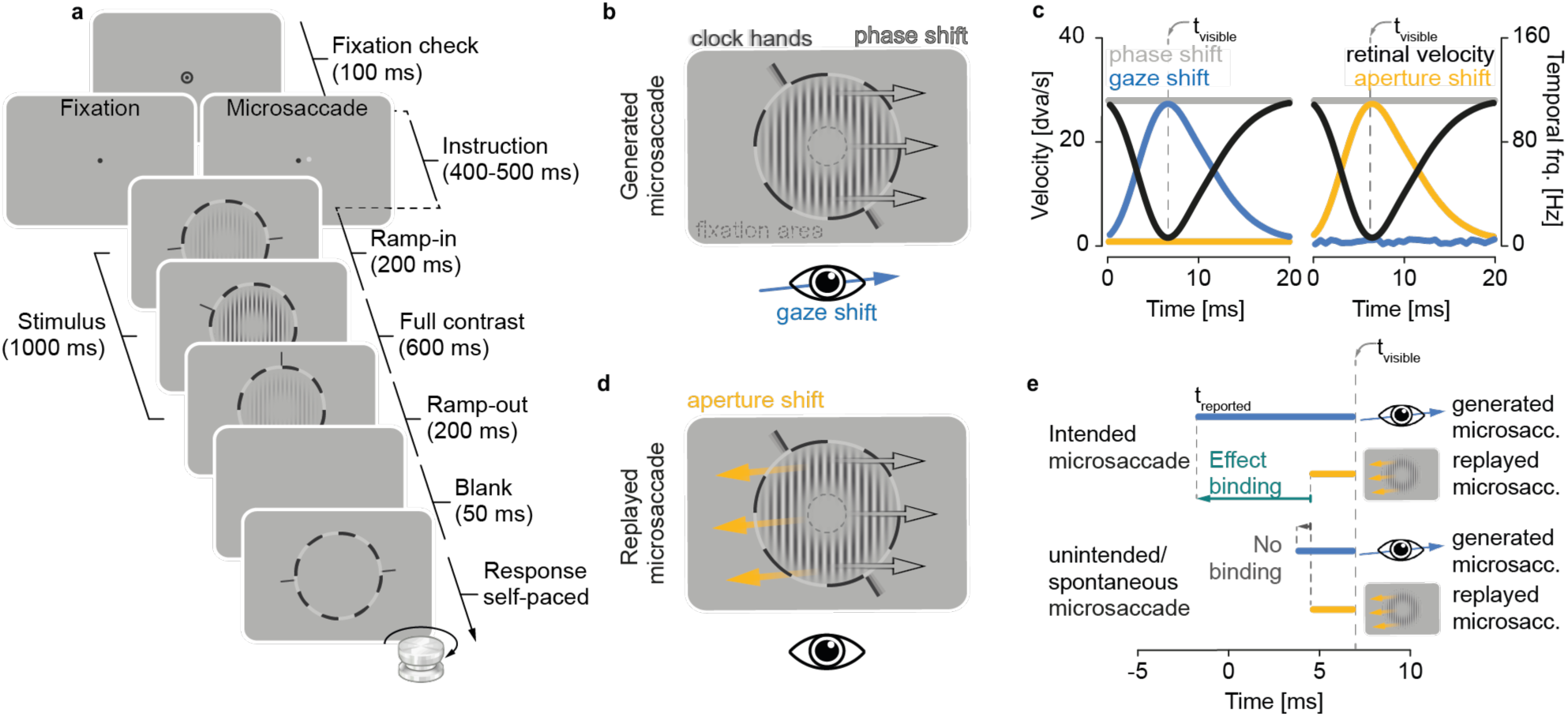
Experimental protocol and stimulus design. **a** Participants were required to pass an initial period of 100 ms of fixation before we displayed eye movement instructions for 400-500 ms: Presentation of a single black dot indicated a fixation trial (in which saccades were labelled as ‘unintended’) and presentation of a black and white dot indicated a microsaccade trial (labelled ‘intended’). In the following, the stimulus was presented together with a clock for 1000 ms. To avoid sudden visual transients, (only) we modulated stimulus contrast with an initial fade-in and final fade-out period (200 ms each). Clock hands moved at a constant speed of 180°/s. After a short blank period of 50 ms, the clock reappeared, and participants were required to adjust the clock hands via rotation knob until they matched their position when the stimulus was visible with pressing the spacebar indicating stimulus absence. Procedure in **Experiment 2** was identical without instruction interval. **b-d** Stimulus display across conditions: If an eye movement was generated in the same direction (b) and with comparable peak velocity (c, left panel, blue curve) as the grating’s rapid temporal phase shift (c, left panel, gray curve), the retinal velocity of the stimulus came close to 0, rendering the stimulus visible for a short period of time (c, left panel, t_visible_ of black curve). An aperture shift in the opposite direction of the phase shift (d) and with the same velocity profile of a microsaccade (c, right panel, yellow curve) similarly rendered the stimulus visible as a short distinct impression of a grating (c, right panel, t_visible_ of black curve). **e** Predictions about effect binding in different conditions: Intended microsaccades should lead to a stronger predating compared a replay of their retinal consequences (relative to the objective time of stimulus visibility t_visible_), with 0 millseconds indicating saccade onset. Temporal estimates should not differ between stimulus conditions for unintended or spontaneous microsaccades.

To measure vFoA and quantify its strength, we relied on the intentional binding effect (Haggard & Clarke, 2003; Haggard 2005; Haggard 2017). The intentional binding effect presumes that the time between an action and its effect is experienced as compressed when FoA is experienced. This effect is bi-directional: Agents experience actions later (‘action binding’) and their effects earlier in time (‘effect binding’) compared to their objective timing. Here, we focused on effect binding both because it is the stronger temporal effect (Haggard, 2017) and because microsaccades are themselves hardly experienced (Klanke et al., 2024)— potentially leading to a high variance in reports if we asked participants when the action was perceived (and not the perceptual consequence). Because effect binding quantifies vFoA using temporal estimates, we presented a clock along with the stimulus and had participants report the time at which they perceived the grating flash. We hypothesized that observers experienced effect binding if they reliably reported stimulus perception earlier when the stimulus was stabilized by a microsaccade compared to its replay (**Fig. 1e**).

Utilizing microsaccades offers several key advantages for investigating FoA compared to other types of actions—including other types of eye movements. Their small size and the strong correlation between amplitude and peak velocity grant significant control over their visual consequences, allowing for a fine-grained classification of each saccade’s potential to be accompanied by vFoA. Moreover, while spontaneous microsaccades occur with limited awareness, intended and unintended microsaccades are consciously experienced (Klanke et al., 2024), facilitating specific and contrasting predictions regarding the expected level of vFoA based on sensorimotor awareness. Especially movement intention is classically seen as a crucial prerequisite for FoA, as unintended movements (e.g., passive limb displacements; Moore et al., 2009) or movements induced by transcranial magnetic stimulation (TMS; Haggard & Clark, 2003) typically lack FoA. This notion, however, is not uncontroversial (Legaspi & Toyoizumi, 2019) and positive evidence for the relevance of intention for FoA altogether rare, as intention is difficult to isolate from movement execution. To test the role of intention for vFoA, therefore, our study directly compared effect binding for three different types of eye movements: intended, unintended (**Experiment 1**), and spontaneous microsaccades (**Experiment 2**). In **Experiment 1**, observers were instructed to execute a small, deliberate saccade to a memorized target location (Willeke et al., 2019) as soon as the fixation point (and saccade target) disappeared (instructed saccade trials; **Fig. 1a**), or to maintain fixation (instructed fixation trials; **Fig. 1a**). We labelled saccades in instructed saccade trials *intended microsaccades* and saccades in instructed fixation trials *unintended microsaccades*. In **Experiment 2**, observers were informed about the spontaneous generation of microsaccades but did not receive specific instructions regarding their microsaccades (**Fig. 1a**). Eye movements generated in this experiment were labelled *spontaneous microsaccades*. By directly comparing intended, unintended and spontaneous microsaccades, we expected to find a vFoA for intended microsaccades, but not for unintended ones (despite observers’ awareness of the eye movements themselves), and equally vFoA for the unconscious spontaneous microsaccades.

## Results

### Motor control for microsaccades

In **Experiment 1**, observers generated unintended microsaccades in a significantly smaller proportion of trials compared to intended ones (*t* (9) = 3.83, *p* = 0.004; unintended: mean = 0.12±0.08; intended: mean = 0.52±0.22; **Fig. 2a**). The number of intended microsaccades was affected by the distance between fixation dot and saccade target (range: 0.2 dva to 1 dva), with larger target distances leading to a higher proportion of trials with microsaccade (one-way repeated-measures analysis of variance, rmANOVA: *F* (4,36) = 11.00, *p* < 0.001; **Fig. 2a**). The proportion of trials with spontaneous microsaccades in **Experiment 2** was in between and significantly different from both intended (*t* (14.7) = 2.23, *p* = 0.042) as well as unintended ones (*t* (14.6) = –2.29, *p* = 0.038; spontaneous = 0.27±0.13). In line with previous studies (Klanke et al., 2024; Hafed & Goffart, 2020; Poletti et al., 2020; Willeke et al., 2019; Rucci & Poletti, 2015; Ko et al., 2010), our data suggest that observers can control microsaccade generation: While unintended microsaccades occurred occasionally, they were generated at a lower rate than even spontaneous microsaccades. Unsurprisingly, observers generated the most microsaccades when instructed to.

**Fig 2.**
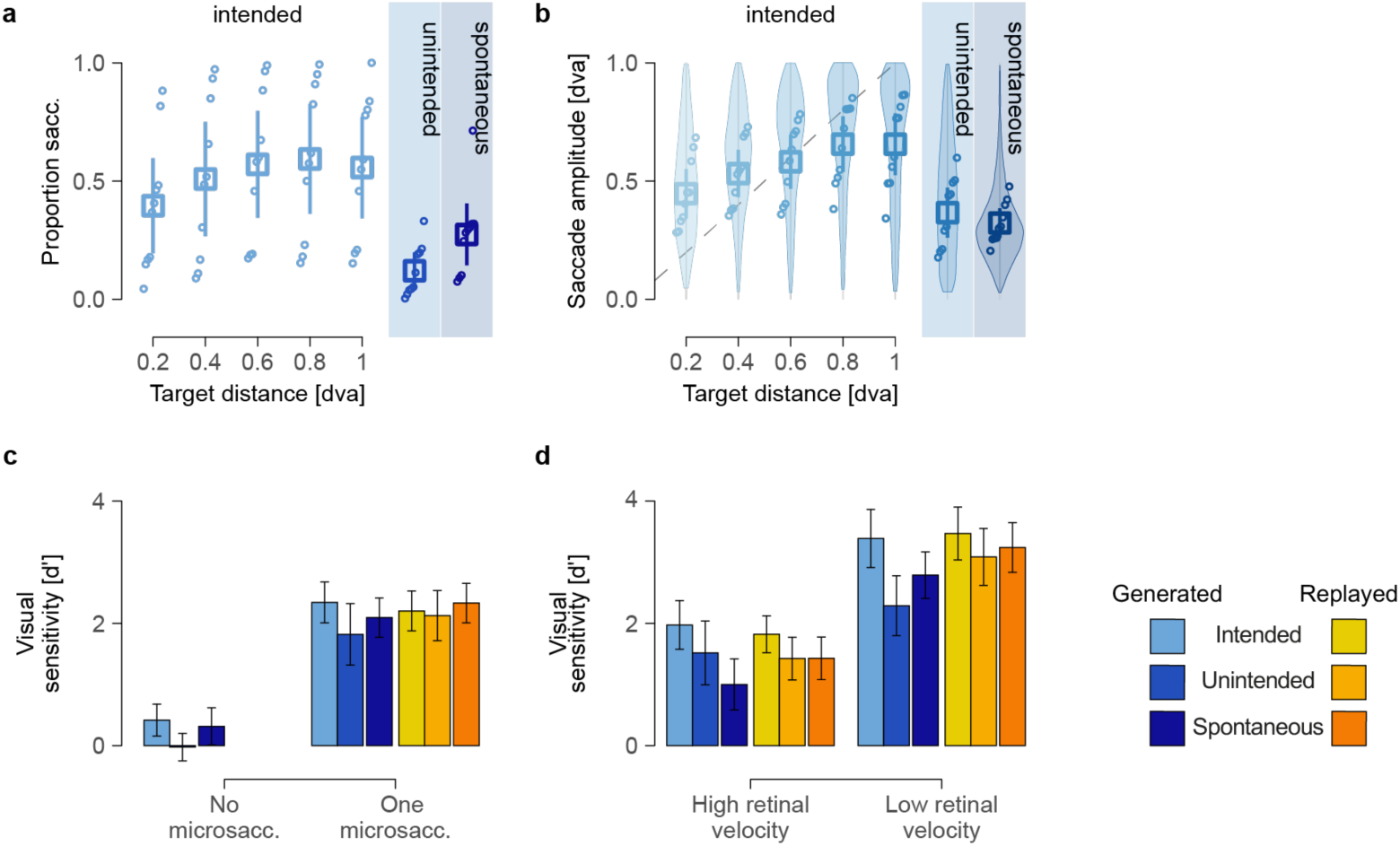
Accurate control of and high visual sensitivity for microsaccades. **a** Proportion of trials of different types of microsaccades and, for intended microsaccades, across target distances ranging from 0.2 to 1 dva. Empty circles represent the average rates for each participant and condition, arranged from lowest to highest values. Squares indicate the group means, and error bars show the 95% confidence intervals. **b** Amplitudes of different types of microsaccades and, for intended microsaccades, across target distances. Empty dots represent the average amplitudes for each participant and target distance, while violin plots illustrate the distribution of all saccade amplitudes. **c** Visual sensitivity as a function of microsaccade generation, across different stimulus conditions (active vs replay) and eye movement types (intended, unintended, and spontaneous). **d** Visual sensitivity as a function of retinal velocity, with low-velocity trials defined by the saccade’s peak velocity being within ±30 dva/s of the grating’s velocity, and high-velocity trials where the saccade’s peak velocity falls outside this range.

Similarly, microsaccade amplitudes were significantly smaller for unintended, compared to intended microsaccades in **Experiment 1** (*t* (9) = 3.40, *p* = 0.008; unintended: mean = 0.37±0.11 dva; intended: mean = 0.57±0.10 dva; **Fig. 2b**). Amplitudes of intended microsaccades increased monotonically with target distance (*F* (4,36) = 30.36, p < 0.001; **Fig. 2b**). A linear mixed effects model, fit to the amplitudes of those intended eye movements, revealed significant positive estimates for all successive difference contrasts (all *p*s < 0.001, except 0.8 vs. 1.0 dva: *t* (4983.4) = –1.27, *p* = 0.204, beta = –0.01±0.02). Amplitudes of spontaneous microsaccades from **Experiment 2** (spontaneous: mean = 0.32±0.06 dva) were smaller than both types of eye movements from **Experiment 1** but statistically only significantly different from intended microsaccades (*t* (14.6) = 4.63, *p* < 0.001). These results suggests that observers successfully adapted their microsaccade amplitudes to the target distances.

### Visual sensitivity to intra-saccadic stimulation

In **Experiment 1**, visual sensitivity was low when no microsaccade was generated (intended: d’ = 0.42±0.30; unintended: d’ = –0.02±0.26) and just failed to reach significance between eye movement types (*t* (9) = 2.21, *p* = 0.054; **Fig. 2c**). Observers were equally insensitive to the stimulus in the absence of spontaneous microsaccades in **Experiment 2** (spontaneous: d’ = 0.31±0.35; **Exp. 1 vs 2**: *t* (12.8) = 0.68, *p* > 0.250; **Fig. 2c**).

Stimulus sensitivity was substantially higher in trials with a microsaccade, irrespective of whether the eye movements were actively generated (d’ = 2.08±0.46) or its visual consequence replayed (d’ = 2.16±0.41) in **Experiment 1**. A two-way rmANOVA revealed no significant difference between stimulus conditions (active vs. replay: *F* (1,9) = 0.18, p > 0.250; **Fig. 2c**). Eye movement type had a significant effect on visual sensitivity (*F* (1,9) = 7.79, *p* = 0.021), with higher sensitivity for intended (d’ = 2.27±0.35) compared to unintended (d’ = 1.97±0.43) microsaccades. Additionally, a significant interaction between stimulus conditions and eye movement type (*F* (1,9) = 6.54, *p* = 0.031) indicated that the difference in sensitivity between eye movement types was more pronounced for generated (*t* (9) = 3.20, *p* = 0.011), compared to replayed eye movements (*t* (9) = 0.71, *p* > 0.250; **Fig. 2c**). In **Experiment 2**, visual sensitivity was similarly high for trials with a generated (d’ = 2.09±0.37) or a replayed (d’ = 2.33±0.37) spontaneous microsaccade. A two-way rmANOVA showed no significant effect of stimulus condition (*F* (1,18) = 2.46, *p* = 0.134) and confirmed that this comparatively high stimulus visibility for trials with a microsaccade (generated or replayed) was consistent across both experiments (**Exp. 1** vs **2**: *F* (1,18) = 0.15, *p* > 0.250). The interaction of experiment and stimulus condition was similarly not significant (*F* (1,18) = 0.56, *p* > 0.250).

Additionally, a match between microsaccade kinematics and stimulus parameters had a considerable impact on visual sensitivity. A three-way rmANOVA for the eye movements from **Experiment 1** demonstrated that matching eye movement and stimulus parameters— inducing retinal speeds below ±30 dva/s—led to a statistically highly significant increase in visual sensitivity (d’ = 3.05±0.47) compared to trials with higher retinal velocities (d’ = 1.68±0.40; *F* (1,8) = 264.24, *p* < 0.001; **Fig. 2d**). While this increase held for both microsaccade types, a significant effect of eye movement type (*F* (1,8) = 18.77, *p* = 0.003) suggests an overall higher sensitivity for intended (d’ = 2.66±0.40) compared to unintended microsaccades (d’ = 2.03±0.47; **Fig. 2d**). The effect of stimulus condition remained insignificant (*F* (1,8) = 1.50, *p* > 0.250), suggesting a comparable sensitivity for replayed (d’ = 2.45±0.41) and generated microsaccades (d’ = 2.22±0.50). However, a significant interaction between microsaccade type and stimulus conditions (*F* (1,8) = 5.84, *p* = 0.042) again indicated that the advantage in visual sensitivity for replayed over generated microsaccades was more pronounced for unintended compared to intended microsaccades. A significant interaction between stimulus condition and retinal velocity (*F* (1,8) = 14.26, *p* = 0.005) confirmed that the difference in sensitivity between generated and replayed microsaccades was more pronounced when retinal velocities were low. This effect was further supported by a significant three-way interaction (*F* (1,8) = 7.2, *p* = 0.028), suggesting that the velocity-dependent differences between replayed and generated microsaccades varied depending on whether the microsaccade was intended or unintended. The remaining interaction between eye movement type and retinal velocity was not significant (*F* (1,8) = 2.76, *p* = 0.135).

A two-way rmANOVA for spontaneous microsaccades from **Experiment 2** revealed an equally highly significant increase in visual sensitivity (*F* (1,9) = 167.44, *p* < 0.001; **Fig. 2d**) for eye movements leading to low (d’ = 3.01±0.43) compared to high (d’ = 1.21±0.41) retinal stimulus velocities. A significant effect of stimulus condition (*F* (1,9) = 17.18, *p* = 0.003) in the absence of an interaction (*F* (1,9) = 0.02, *p* > 0.250) indicated that, similarly to unintended eye movements from the first experiment, visual sensitivity is increased for replayed (d’ = 2.33±041 spontaneous microsaccades compared to generated ones (d’ = 1.89±0.41; **Fig. 2d**). Comparing visual sensitivity between eye movements from the two experiments demonstrates no significant differences overall (**Exp. 1 vs 2:** *t* (16.7) = –0.86, *p* > 0.250).

Our analysis of visual sensitivity demonstrated that our high-temporal frequency stimulus was invisible during stable fixation but became visible when a microsaccade was generated or replayed. High sensitivity when eye movement and stimulus parameters matched, points towards gaze-contingent retinal stabilization as the most important contributor for stimulus perception. Increased sensitivity for intended compared to unintended microsaccades can be attributed to their larger amplitudes and, concurrently, higher peak velocities (see SUPPLEMENTARY MATERIAL *Parameters of different types of generated and replayed microsaccades*) leading to an overall better match of parameters and, hence, greater degree of retinal stabilization. This reduced retinal stabilization for unintended microsaccades resulted in fewer low-retinal velocity trials and consequently a smaller and more variable sample. Importantly, microsaccade parameters were generally well matched across stimulus conditions, supporting the validity of the comparison between generated and replayed microsaccades. Observers’ higher sensitivity to replayed microsaccades might be due to an overestimation of saccade peak velocities of the video-based eye tracker (cf. Schweitzer & Rolfs, 2022), which would have led to a higher peak velocity for replays than for generated saccades, and, as a consequence, a lower retinal velocity of the stimulus. Overall, we conclude that our stimulus and paradigm worked as intended and the stimulus perception was well matched between stimulus conditions and eye movement types.

### Feeling of agency for immediate visual consequences of eye movements

To determine whether observers experience vFoA for microsaccades, we compared temporal estimates for generated and replayed eye movements. A tendency to report stimulus perception earlier in trials with generated microsaccades (i.e., active condition), compared to its timing when the retinal consequences were replayed (i.e., replay condition), would suggest effect binding (cf. **Fig. 1e**) and thereby indicate vFoA, consistent with the intentional binding effect.

We used linear mixed-effects models (LMEs) to predict temporal estimates as a function of stimulus condition (active vs. replayed microsaccade) for each eye movement type (intended, unintended, spontaneous). For intended microsaccades from **Experiment 1**, the LME yielded a positive effect of stimulus condition (estimate = 3.34 ms), indicating a stronger predating of stimulus percept for generated microsaccades (mean = 41.98±74.45 ms) compared to replayed microsaccades (mean = 45.32±54.29 ms). While this difference was in the predicted direction, the effect was not statistically significant (*t* (7.8) = 0.20, *p* > 0.250; **Fig. 3a**, panel 1), providing no evidence in support of effect binding for intended microsaccades. To further evaluate the absence of an effect, we conducted a Bayesian analysis, which yielded strong evidence against the hypothesis that stimulus condition significantly influenced the timing of stimulus visibility (intended: BF_10_ = 0.07). We repeated the same analysis for unintended microsaccades from **Experiment 1**. The LME yielded an estimate very close to zero (estimate = 1.21 ms), indicating that the subjective timing of stimulus perception reported by observers was nearly identical between the active condition (mean = 30.31±57.91 ms) and the replay condition (mean = 31.52±50.11 ms). Additionally, the result for unintended microsaccades was statistically insignificant (*t* (4.9) = 0.06, *p* > 0.250; **Fig. 3a**, panel 2), indicating an absence of effect binding for unintended microsaccades. The Bayesian analysis for unintended microsaccades also yielded strong evidence against the hypothesis of a significant stimulus condition effect (unintended: BF_10_ = 0.08).

**Fig 3.**
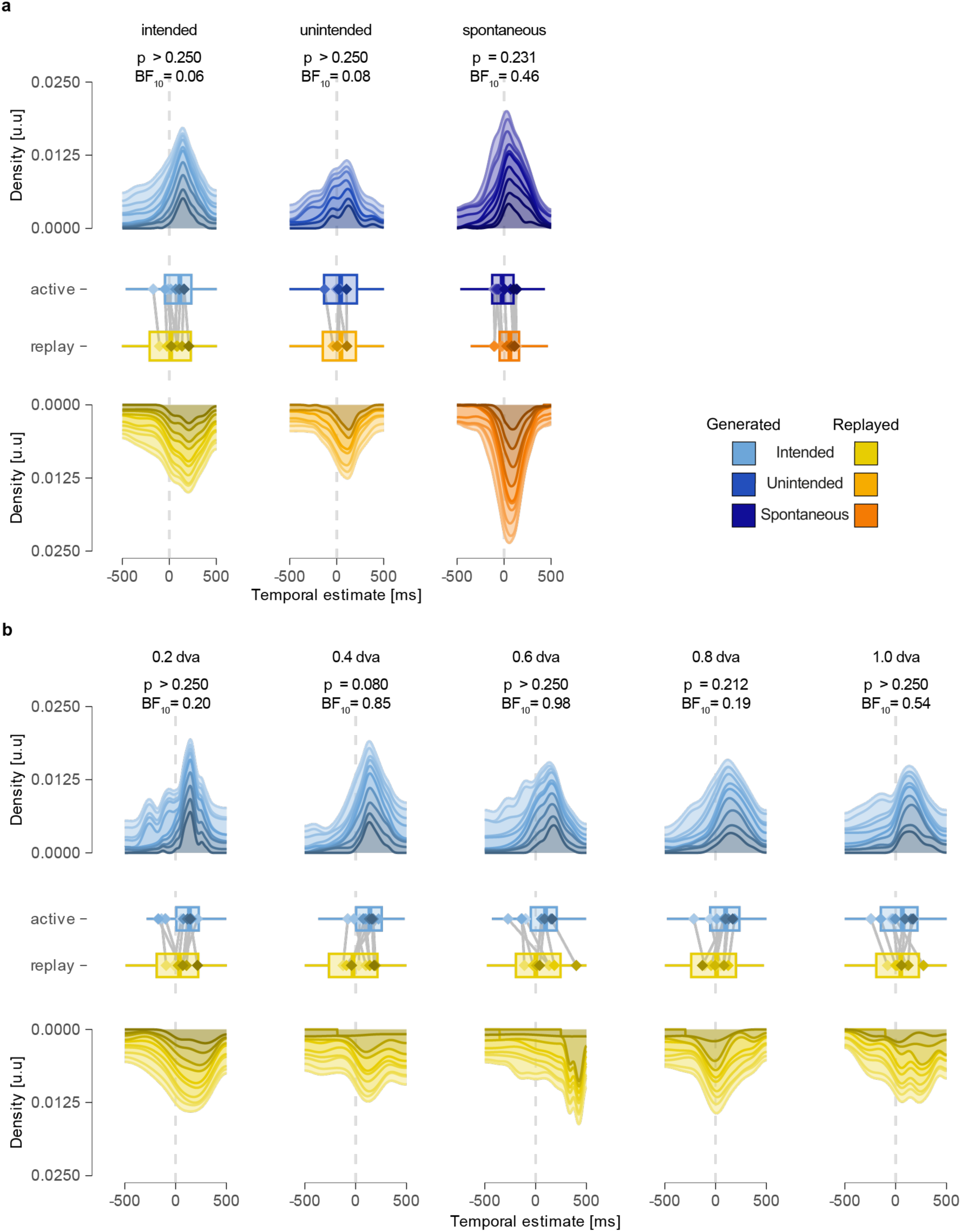
Absence of a conscious feeling of agency for intended, unintended, and spontaneous microsaccades. **a** The upper and lower rows show the stacked density of time intervals (report–event) between the time of the highest retinal velocity of the stimulus (event) and the reported time of the stimulus (report) in the active (blue) and replay (yellow) conditions for different eye movement types. Different shades represent data from individual participants. The box plots in the middle panel depict the median, interquartile range, and overall spread for each condition and eye movement type, with outliers omitted for clarity. Diamond shapes indicate distribution means for individual participants (with colors corresponding between diamond and density per participant), while vertical lines connect values for each participant to illustrate the direction of the effect. **b** This panel presents the same measures as in a, but for intended eye movements only with separate target distances (ranging from 0.2–1.0 dva) in subpanels. All other aspects are the same as in a.

For spontaneous microsaccades from **Experiment 2**, we found a positive yet insignificant estimate for the effect of stimulus condition (estimate = 11.78 ms; *t* (7.6) = 1.30, *p* = 0.231; **Fig. 3a**, panel 3). Consistent with the positive estimate, our data showed a similar trend to that observed for intended microsaccades, with earlier subjective reports when the stimulus was perceived as a consequence of a generated microsaccade (mean = 17.82±64.08 ms) compared to a replayed microsaccade (mean = 29.60±63.49 ms). While the Bayes factor was slightly higher for spontaneous microsaccades (BF_10_ = 0.46), it still provided moderate evidence against the hypothesis that the stimulus condition significantly impacted the time of stimulus visibility.

Finally, to determine if effect binding of intended microsaccades was affected by the amplitude of an eye movement, we used linear mixed-effects models (LMEs) to predict temporal estimates of stimulus perception as a function of stimulus condition (active vs. replayed), separately for each target distance (ranging from 0.2 to 1 dva). We used target distance as a discrete predictor because microsaccade amplitudes tend to match target amplitudes and hence covary with target distance (see Results section, Motor control for microsaccades). Additionally, we performed Bayesian analyses to evaluate the evidence for or against the hypothesis that temporal estimates differed between active and replayed conditions for intended microsaccades across target distances. We found little evidence for a significant effect (all *p*-values ≥ 0.080) or support of the test hypothesis (BF_10_ ≤ 0.99) for all five target distances (see **Fig. 3b**, panels 1-5).

Together, these results demonstrate the absence of effect binding across all types of eye movements tested here and, thus, a lack of evidence for the hypothesis that participants experienced vFoA for either intended or unintended microsaccades from **Experiment 1**, nor spontaneous ones from **Experiment 2**.

## Discussion

### No effect binding for the immediate sensory consequences of minuscule eye movements

We investigated the feeling of control for minuscule eye movements and the changes in visual information they bring. We found no evidence for a predating of the sensory consequence of a microsaccade compared to the perceived timing of the visually identical yet purely perceptual replay of retinal consequences characteristic for effect binding: Reported timing of stimulus perception did not differ significantly between generated and replayed eye movements for intended or unintended microsaccades (**Exp. 1,** cf. **Fig. 3a**, panels 1-2), nor did we find such a difference for spontaneous microsaccades (**Exp. 2**, panel 3). The absence of effect binding in our data—at the very least—does not support the idea of vFoA: A feeling of control over the immediate sensory consequences of microsaccades. The consistent absence of effect binding for intended microsaccades (**Exp. 1**), irrespective of target distance, additionally suggests that intended eye movements are not automatically accompanied by a sense of control of the immediate visual consequence and intention is, therefore, not sufficient for FoA in general and vFoA in particular (cf. **Fig 3b** panels 1-5; see *Feeling of agency for immediate visual consequences of eye movements* for more details). To corroborate these results and investigate the amount of evidence against the test hypothesis, we calculated Bayes factors across all eye movement types. Evidence in favor of the null hypothesis consistently outweighed evidence for the test hypothesis—particularly for intended microsaccades, the type of eye movement for which we had predicted the strongest effect binding (see *Feeling of agency for immediate visual consequences of eye movements*).

### No vFoA for the sensory consequences of minuscule eye movements?

One potential conclusion to be drawn from the data presented here is that the absence of effect binding for minuscule eye movements implies that there is no vFoA and, hence, no feeling of control for the immediate visual consequences of eye movements. While it may sound paradoxical—normally, we do not experience our eye movements as uncontrolled—it is entirely possible that our feeling of control does not include the *immediate* visual consequences of an eye movement. We consider it possible that observers can experience agency for eye movements in the same way they do for other types of physical actions: Directing one’s gaze at a ‘gaze-activated’ light switch, followed by the light turning on, might well be accompanied by a sense of agency, as classically defined. We however, examined FoA for the changes of the visual scene inherent to every eye movement and, hence, a much less specific category of action. The absence of vFoA, if understood as genuine, might hence indicate that the feeling of control we normally experience for eye movements does not extend to their visual consequences and is bound to the eye movement alone. A way to measure this would be to focus on action binding, a perceived shift in time of the action towards its consequence. While we did not investigate action binding, tentative evidence for this idea can be derived by the fact that our observers generally overestimated the timing of events—and most strongly in the intended microsaccade condition (cf. **Fig 3a**). Since the timing of action and effect is identical in our paradigm (as the visual consequence of a microsaccade is visible as the microsaccade is generated), the overestimation of the event might indicate that the perceived timing of the action is shifted forward in time. The absence of vFoA might also be attributed to microsaccades (alone), as eye movements that are too small, too automatic, or too faint to elicit vFoA. Observers can, however, reliably detect intended and even unintended microsaccades consciously—despite their small size and ephemeral nature (Klanke et al., 2024). Nevertheless, we failed to measure effect binding even for intended microsaccades (cf. **Fig. 3a**, panel 1), which are instances where microsaccades are generated consciously and align with a preceding intention. These findings suggest that microsaccades serve as prototypical eye movements, that allow for more general conclusions.

An alternative interpretation of our findings suggests that vFoA is real, but the immediate sensory consequences of eye movements are not shifted forward in time (i.e., visual consequences are not in the same way bound to their causes as other bodily movements). A vast body of literature calls into question the link between intentional binding and FoA. Evidence exists for intentional binding occurring in the absence of intention or even an action (Suzuki et al., 2019), as well as for unintended actions (Ruess et al., 2020b)—suggesting that intentional binding is neither necessary nor sufficient for effect binding. Though an open debate, intentional binding may be an expression of the expectancy of the consequence of an action (or other type of strong event-effect association) rather than FoA itself (Ruess et al., 2020a). Despite these examples of doubt of intentional binding as an indicator for FoA, more than 20 years of research has demonstrated a connection between the perceived time of an action and effect and the feeling of control over it (Haggard, 2017; Moore & Obhi, 2012), with some studies showing that different measures of agency—including intentional binding— correlate (Imaizumi & Tanno, 2019). Together, this literature lends credence to intentional binding as a measure of agency. We are confident, therefore that the absence of effect binding, as measured by our experiments, implies an absence of vFoA for minuscule eye movements.

### (Re-) Evaluating the paradigm

We maintain that our paradigm worked as intended: Observers generated intended microsaccades when instructed to saccade and largely suppressed unintended eye movements when instructed to fixate (**Exp.1**), with an intermediate rate of spontaneous microsaccades (**Exp. 2**) that was significantly different from both other eye movement types (cf. **Fig. 2a**). Amplitudes of intended microsaccade (**Exp. 1**) were significantly larger than unintended (**Exp. 1**) and spontaneous eye movements (**Exp. 2**). Increasing amplitudes over larger target distances for intended microsaccades additionally reflecting the deliberate modulation of eye movement dynamics in response to task demands as well as a fine-grained motor control (cf. **Fig. 2b**; see *Motor control for microsaccades for details*). Our data on visual sensitivity support the same conclusion: Observers’ visual sensitivity was around zero when no microsaccade was generated but increased significantly in trials with a generated or replayed eye movement—irrespective of microsaccade type (cf. **Fig. 2c**). Significantly higher stimulus sensitivity in trials with matching microsaccade and stimulus parameters—leading to lower retinal stimulus velocity and, hence, a more stable image of the stimulus on the retina. This was true for intended and unintended (**Exp 1**) as well as spontaneous (**Exp. 2**) microsaccades. Moreover, stimulus visibility was well matched between active and replay condition trials for all microsaccades, suggesting a close match of the perceptual consequence of an eye movement and the replay of its retinal consequences (cd. **Fig. 2d**; see also *Visual sensitivity to intra-saccadic stimulation* for details). It is worth noting that both findings on saccade generation and visual sensitivity closely match the ones reported by Klanke et al., (2024), who used a similar stimulus and experimental design.

Nevertheless, a potential reason for the lack of evidence for vFoA in our data might, of course, be that our paradigm failed to allow us to measure it. Two specific limitations warrant consideration: A side effect of how we created a microsaccade-contingent stimulus is that an eye movement and its visual consequences are not necessarily strongly linked: Microsaccades were neither necessary (cf. replayed microsaccades) nor sufficient (cf. no-stimulus condition) to render the stimulus visible. Stimulus perception, on the other hand, provided equally little information about the preceding eye movement behavior: A replayed microsaccade could render the stimulus visible even without an active eye movement, while the stimulus remained invisible in the no-stimulus condition or when the stimulus and microsaccade parameters were too disparate for retinal stabilization. It might, hence, be that we failed to measure effect binding because the correlation between microsaccade and sensory consequence was insufficient to induce a strong feeling of control over the action-effect contingency.

A second reason for why our paradigm might have failed to capture vFoA is similar yet slightly different: While the stimulus was saccade-contingent, it hardly resembled how we experience saccade-contingent changes in the retinal image in everyday life: Our stimulus was perceived during the saccade while pre- and post-saccadic retinal images were largely identical. This contrasts with natural situations, in which the retinal input changes drastically across saccades. If we assume a genuine experience of control over the visual consequences of our eye movements, it seems more plausible that observers perceive control over the targeted change from the pre- to post-saccadic image (e.g., the impression of an apple in the periphery transitioning to the impression of a foveated apple) rather than for an image that is visible only during the saccade. Indeed, visual information received around the time of the saccade routinely escapes conscious perception (c.f., saccadic omission; Matin 1975; Campbell & Wurtz, 1978; Schweitzer et al., 2025). This discrepancy between the gaze-contingent stimulus in our task and the natural perceptual consequences of saccades may explain why our paradigm failed to provide evidence supporting vFoA for minuscule eye movements, despite the high degree of control we experience over how our gaze shifts through the visual scene in everyday life.

### Eye movements time perception, and the chronostasis illusion

An alternative explanation for the absence of effect binding in our data (that still maintains the possibility of vFoA) is that temporal processing around saccades is poor (Morrone et al., 2005), and any potential binding effect may have been obscured by the observers’ coarse ability to process time before, during, and after an eye movement. This is likely the case for at least some of the participants in **Experiment 1**, as the distribution of temporal estimates is fairly spread out and, for some, almost uniform (cf. **Fig. 3a**, panels 1 and 2). **Experiment 1** involved a dual-task paradigm, combining instructed eye movement behavior with the time estimation task. This likely diminished observers’ ability to track the positions of the clock hands (Schwarz & Weller, 2023), as dual-task paradigms are frequently—if not always—characterized by a division of attention between tasks, often resulting in performance declines in one or both tasks (Pashler, 1994). In **Experiment 2**, participants were only instructed to report the timing of the stimulus flash, while all microsaccades occurred spontaneously and without any instruction. This allowed participants to fully focus on the time estimation task, leading to more precise estimates for almost all participants (cf. **Fig. 3a**, panel 3).

Nevertheless, temporal processing deficits around saccades do not necessarily prevent measuring temporal effects. One such effect, that is particularly interesting to consider in the context effect binding and vFoA is an illusion known as the *chronostasis effect*: A perceived freezing of time (in fact; the movement of the clock hand) after making a saccade towards a clock face (Brown & Rothwell, 1997). The duration of the post-saccadic freezing reportedly corresponds to the saccade duration (Yarrow et al., 2001, 2006). Similar findings of a pre-dating of the post-saccadic percept to before the new fixation (Hunt & Cavanagh, 2009) support this idea. Nevertheless, it stands to reason that effect binding of the post-saccadic retinal image is characteristic for vFoA as the consequence of an eye movement can serve as an equally viable explanation for the chronostasis illusion and similar temporal effects.

## Conclusion

In this study, we examined the subjective feeling of control over an eye movement and its visual consequence: vFoA. Across two experiments, we found no evidence for effect binding— a temporal shift in perceived stimulus time—as a marker of agency for these minute eye movements. Perceived timing of visual stimuli did not differ between self-generated and replayed microsaccades, regardless of whether the movements were intended, unintended (**Exp. 1**), or spontaneous (**Exp. 2**). These findings suggest that the mere generation of a microsaccade, even when intended, does not inherently produce a sense of control over its visual outcome. Thus, to the extent that effect binding is a reliable consequence of vFOA, our results challenge that vFoA arises automatically with intentional microsaccades. Thus, intention alone may not be sufficient to generate a feeling of agency for the immediate visual consequences of eye movements.

## Acknowledgements

JNK was supported by the Berlin School of Mind and Brain, Humboldt-Universität zu Berlin. SO was supported by the DFG (OH 274/4-1) and funding from the Heisenberg Programme of the DFG (OH 274/5-1). MR was supported by the European Research Council (ERC) under the EU’s Horizon 2020 research and innovation program (grant agreement no. 865715), and by the DFG (grants RO3579/8-1 and RO3579/10-1).

## STAR Methods

### RESOURCE AVAILABILITY

#### Lead contact

Information and requests regarding resources for this study should be directed to and will be fulfilled by the lead contact, Jan-Nikolas Klanke

#### Materials availability

There are no restrictions for the distribution of materials.

#### Data and code availability

- The preregistration, data, and all original code for **Experiment 1** has been deposited at the Open Science Framework and will be made publicly available as of the date of publication. [LINK WILL FOLLOW HERE UPON PUBLICATION].
- The preregistration, data, and all original code for **Experiment 2** has been deposited at the Open Science Framework and will be made publicly available as of the date of publication. [LINK WILL FOLLOW HERE UPON PUBLICATION].

### EXPERIMENTAL MODEL AND SUBJECT DETAILS

In **Experiment 1**, a total of 10 participants were recruited by means of the “Psychologischer Experimental-Server Adlershof” (PESA) of the Humboldt-Universität zu Berlin. Participants (6 female, 1 diverse) had a mean age of 28.0 years old (*SD* = 4.9, *min* = 22, *max* = 38), and all 10 were right-handed and 7 were right-eye dominant. All 10 participants had normal or corrected-to-normal vision. Participants were paid upon completion of the last session. The compensation was based on an hourly rate of €10/hour. Alternatively, psychology students could choose to obtain participation credit (1 credit per 15 minutes of participation) required for the successful completion of their bachelors’ program.

In **Experiment 2**, we recruited a total of 10 participants. We again recruited participants by means of the “Psychologischer Experimental-Server Adlershof” (PESA) of the Humboldt-Universität zu Berlin. Participants (7 female, 0 diverse) had a mean age of 26.7 years old (*SD* = 7.8, *min* = 21, *max* = 34). Nine participants were right-handed, and six right-eye dominant. All participants had normal or corrected-to-normal vision. Participants were paid upon completion of each experiment: Compensation was based on an hourly rate of €8/hour and a bonus payment of €4 for the completion of the final session. Psychology students could again alternatively choose to gain participation credit (1 credit per 15 minutes of participation) required for the successful completion of their bachelors’ program.

**Experiment 1** and **2** were approved by the ethics committee (Ethikkomission) of the Institut für Psychologie at the Humboldt-Universität zu Berlin and conducted in agreement with the Declaration of Helsinki (‘World Medical Association Declaration of Helsinki’, 2013) and the General Data Protection Regulation (GDPR) of the EU. All participants provided informed consent in writing before the start of the first session.

For **Experiment 1** and **2**, we pre-registered three exclusion criteria that ensured that participants would not participate if they showed the inability to execute stable fixation and correct eye movements:

- The inability to complete at least 4 blocks during the first experimental session due to fixation failures led to immediate exclusion from the experiment.
- If we could not detect more than 2.5 (**Exp. 1**) or 2.5 microsaccades (**Exp. 2**) in the crucial time window (200–800 ms relative to stimulus onset) of trials with generated eye movements across each block of the first session, we likewise excluded the participant from further testing.
- During data analysis, we double-checked the eye movement data offline. We excluded all participants that generated less than 150 microsaccades in the crucial time window (200–800 ms relative to stimulus onset) of generated microsaccade condition trials. Especially the second and third criteria were set to ensure that we would obtain enough data from each participant for the planned analyses.

In **Experiment 1**, nine participants were excluded from data collection: Five participants were excluded after the first session for creating less than 2.5 microsaccades in the crucial time window (200–800 ms after stimulus onset) of the active condition trials in each block of the first session. One participant decided against further participation for personal reasons. Lastly, we excluded two participants who generated much less than the required 150 microsaccades (70 and 74 respectively). To not bias our data sample too strongly, we kept two participants who generated only slightly less than the required amount (136 and 140 microsaccades respectively). One more participant was excluded after completing four sessions. While they generated more than 15 microsaccades in the first session, the overall extremely low number of microsaccades in the subsequent three sessions made it extremely unlikely that the participant would come close to the required 150 microsaccades (total of 37 microsaccades after four sessions).

In **Experiment 2**, a total of four participants were excluded: one participant was excluded because of their inability to fixate (less than 3 blocks of data in the first session of the experiment). Two participants were excluded because they generated less than the required amount of microsaccades in the first session of the experiment. Finally, one participant was excluded due to an overall low number of microsaccades over the curse of the entire experiment (79 microsaccades). We additionally excluded two sessions of the first participant due to an error in the code that led to an erroneous display of the stimulus.

### METHOD DETAILS

#### Apparatus

Participants were seated in a dark room in front of a screen at a distance of 340 cm and their head stabilized using a chin rest. We projected visual stimuli on a 141.0 x 250.2 cm video-projection screen (Stewart Silver 5D Deluxe; Stewart Filmscreen, Torrance, CA, USA) using a PROPixx DLP (960 × 540 pixels; VPixx Technologies Inc., Saint Bruno, QC, Canada) with a refresh rate of 1440 Hz. We recorded participants’ eye positions of both eyes with a head-mounted eye tracker at a sampling rate of 500 Hz (EyeLink 2 Head Mount; SR Research, Ottawa, ON, Canada). The experiments were controlled on a workstation running the Debian 8 operating system, using Matlab (Mathworks, Natick, MA), the Psychophysics Toolbox 3 (Brainard, 1997; Kleiner et al., 2007; Pelli, 1997) and the EyeLink Toolbox (Cornelissen et al., 2002).

### The stimulus

#### Active condition

We used the same stimulus in **Experiment 1** and **2**: a vertically oriented sinusoidal grating multiplied with a tapered cosine mask. Together, grating and mask created the appearance of a striped donut that smoothly blends in with the background (cf. **Fig. 1b/c**). The stimulus had a total diameter of 10 dva with a tapered section of 1/3, resulting in an inner, circular, opaque section of the stimulus with approximately 3 dva in diameter (dashed line within the inner part of the stimulus in **Fig. 1b/d**). The stimulus was always presented centered on the gaze position of the participant and the mask, therefore, occluded the area of the grating to which participants eye positions were restricted (if eye positions were tracked outside this area, the trial was aborted and repeated in the end of each block). Lastly, the grating had a spatial frequency of 4 cpd and a maximum contrast of 50%. Additionally, phase onset was randomized in each trial.

Our stimulus was designed to be invisible during fixation, but visible when briefly stabilized on the retina by a microsaccades with matching kinematics. To achieve invisibility under stable fixation conditions, a high-velocity phase shift was added to the grating—creating a temporal frequency above 60 Hz (cf. Castet & Masson, 2000). In the first session of the experiment, the speed of the phase shift was fixed to 27.6 dva/s (i.e., 110.4 Hz) for all participants. This value corresponds to a microsaccade with a 0.5 dva amplitude, based on the main sequence function—a known curvilinear relationship between saccade amplitude and peak velocity (Zuber et al., 1965)—and values reported by Collewijn et al. (1988). In later sessions of **Experiment 1,** the phase shift velocity was set to at least 27.6 dva/s but was otherwise adjusted based on the microsaccade parameters from each participant’s first session. To achieve high retinal stability for most, but not all, microsaccades, we fit the main sequence function of Collewijn et al. (1988) to each participant’s microsaccade data and extracted parameters for which 75% of the microsaccades had a higher peak velocity. This individualized phase shift velocity ensured comparable stimulus visibility across participants, despite potential differences in eye movement behavior. In **Experiment 2**, the phase shift velocity was fixed to 27.6 dva/s for all participants and sessions (to maximize visibility).

The direction of the phase shift—determined by the stimulus orientation—was either rightward (stimulus orientation of 0 deg from vertical) or leftward (stimulus orientation of 180 deg from vertical) to match the direction of potential microsaccades (microsaccades are most likely to occur in horizontal directions, see Engbert & Kliegl, 2003). Stimulus presentation started at a contrast of 0 and was ramped up to maximum contrast (of 50%) within 200 ms. Conversely, in the last 200 ms of the presentation, the stimulus’ contrast slowly decreased and the stimulus, hence, fade out. This was done to avoid sudden stimulus on- and offsets that might have led to unintended transient changes in stimulus visibility. Because generated microsaccades rendered it visible, this stimulus condition was called active condition.

#### Replay Condition

In the replay condition, the stimulus was identical to that in the active condition regarding its size, tapered section, orientation, contrast modulation, phase onset, phase shift velocity, as well as its’ initial placement and presentation duration (see **Fig. 1c/d**). It differed in that we added a rapid change in the onscreen location of the stimulus aperture. This was done to replay the retinal consequence of a previous eye movement back to the observer. We modeled the aperture motion based on individual microsaccade data from each participant, following a strict detection and pre-processing pipeline. (see section *Pre-processing* in the **STAR Methods** below).

#### No-stimulus condition

In the no-stimulus condition, the stimulus was presented at 0% contrast while all other parameters were identical to active and replay condition trials. In **Experiments 1** and **2**, 20% of the trials were no-stimulus condition trials, while the active and replay condition trials each comprised 40% of the total. All stimulus conditions were randomly interleaved within each block.

### General Methods

#### Eye movement task

In **Experiment 1**, our goal was to directly compare intended and unintended microsaccades. To achieve this, we used a modified version of the memory-guided microsaccade paradigm (Willeke et al., 2019), which we had previously employed in another study to investigate the role of intention in microsaccade awareness (Klanke et al., 2024). For trials involving unintended microsaccades, we provided explicit instructions for participants to maintain their gaze at a fixed onscreen location. The location was indicated by a black dot with a diameter of 0.2 dva that was presented at the beginning of each trial (50% of all trials). In contrast, for trials where participants were instructed to generate a microsaccade intentionally, a microsaccade target was presented alongside the fixation point: a white dot with a diameter of 0.2 dva (remaining 50% of trials). This target was presented at a radial distance of either 0.2 dva, 0.4 dva, 0.6 dva, 0.8 dva, or 1 dva relative to the fixation dot. To introduce some variation to the microsaccade target location, the target dot was allowed to be displaced along the circumference of its radial distance to the fixation dot. This displacement was sampled from a normal distribution centered on 0 deg and with a standard deviation of 25 deg, thus resulting in onscreen locations of the target with a small vertical offset. Both instruction conditions were presented during the initial fixation interval, after which the fixation dot and saccade target (if present) were no longer displayed. Participants were instructed to maintain fixation on the target location even after it disappeared, and only shift their gaze to the (remembered) location of the microsaccade target once it was no longer visible on the screen.

### Procedure

#### Fixation-check interval

Before the start of each trial, a target-shaped central fixation point appeared before an otherwise grey background. The inner part of the fixation target had a diameter of 0.2 dva while the outer ring had a diameter of 0.6 dva. Before the onset of each trial, we ran a fixation control routine that required the gaze position of the observer to be inside a circular region (3 dva in diameter) around the fixation point. When the fixation control was successful for at least 100 ms, the trial began. The center of the fixation target appeared at the onscreen location on which the stimulus presentation was centered in the following.

#### Fixation interval

The fixation interval (present only in **Experiment 2**) started as soon as the outer ring of the fixation target disappeared. The duration of the fixation interval varied randomly between 400 ms and 500 ms to avoid routine anticipatory eye movements. Fixation dot and microsaccade target remained visible for the entire duration of the fixation interval. Participants were instructed to keep their gaze trained on the fixation dot as long as it was visible (i.e., for the duration of the fixation interval). If a microsaccade target was displayed additionally, participants were to memorize the onscreen location of the target and generate an eye movement to this location as soon as the fixation dot and microsaccade target disappeared (i.e., in the beginning of the stimulus presentation interval). In trials without microsaccade target presentation, participants were instructed to keep their eye position centered on the location of the fixation dot even after it disappeared.

#### Stimulus presentation interval

The disappearance of the fixation point (inner part and outer part) and the appearance of the clock face and hands indicated the start of the stimulus presentation interval. Stimulus presentation lasted for 1000 ms independent of stimulus condition and eye movement instruction. The position of the stimulus was determined randomly in each trial, but its midpoint was always within ±4 dva relative to the screen center (horizontally as well as vertically). Between the stimulus presentation and the response interval, there was a short delay of 50 ms during which nothing was presented on the gray screen.

#### Response interval

At the beginning of the response interval, only the clock face and hands reappeared. The participants’ task was always the same, regardless of the condition: If participants had perceived the stimulus during the trial, they had to report its timing by adjusting the clock hands. In other words, participants had to replicate the positions of the clock hands when the stimulus was visible during the moment of relative retinal stabilization. The adjustment of the clock hands was carried out by rotating a volume knob, and participants had to confirm their selection by pressing a button next to the dial. If participants did not perceive the stimulus, they were instructed to indicate this by pressing the space key. There was no temporal limitation on the response time, meaning the response was self-paced.

#### Variations in Experiment 2

The fixation check, stimulus presentation, and response intervals were the same in **Experiments 1** and **2**. To compare intended and unintended microsaccades with spontaneous ones, we removed the fixation interval (before the stimulus presentation), and no eye movement instructions were presented. Before the first session, participants were informed that microsaccades could occur spontaneously and that trials would abort if their gaze position deviated too much from the location indicated during the fixation check interval. No received no further instruction were administered regarding participants eye movement behavior.

### Online control of eye positions

Throughout **Experiment 1** and **2**, participants’ eye positions were continuously tracked. Eye and screen coordinates were calibrated and validated using a standard nine-point procedure before the first trial of each session and whenever needed. Blinks and gaze deviations greater than 1.5 dva from fixation were also monitored in both experiments, resulting in the termination of the trial. Aborted trials were repeated at the end of each block in a randomized order.

### Pre-processing

Binocular microsaccades were identified using the algorithm outlined by Engbert and Mergenthaler (2006) in both **Experiment 1** and **2**. The velocity threshold was set to a λ of 5, and the minimum microsaccade duration was 6 ms (equivalent to 3 data samples). To avoid including potential over- or undershoot corrections, two microsaccades were combined if the interval between them was less than 10 ms (5 data samples).

The gaze positions of the dominant eye of each observer, recorded during binocularly detected microsaccades, were used for the replay of the retinal consequence of a microsaccade. We only replayed the retinal consequences of microsaccades that were recorded within the crucial time window (200–800 ms after stimulus onset) of trials where a generated microsaccade could render the stimulus visible. This was done to enable a direct comparison of stimulus visibility across conditions. We did not use the raw microsaccade data for the stimulus but pre-processed the recorded gaze trajectories. In a first step, we subtracted the coordinates of the first data sample form all remaining samples and re-centered the recorded gaze positions of each microsaccade on the origin. In a second step, we excluded all gaze positions recorded after the microsaccade reached its maximum amplitude to ensure that microsaccades, that follow a curved trajectory, did not lead to blurry or obscure percepts when replayed. Microsaccades were excluded from being replayed if their duration was shorter than 6 ms (3 data samples) before reaching their maximum amplitude. In a third step, recorded eye positions were projected onto the saccade vector by recalculating the location of each gaze position during the saccade relative to its amplitude. In a fourth step, we fit a gamma function to the velocity profile of this saccade vector and determined optimal fits by means of a root mean square error (RMSE) procedure. We additionally excluded microsaccades for which the RMSE deviated more than two standard deviations from the mean of all RMSEs of the same session from one participant. In step number five, gaze positions were redistributed along the saccade vector based on the fitted velocities. To compensate for the difference between the eye tracker’s frequency (500 Hz) and the projector’s refresh rate (1440 Hz), we combined the recalculation of the gaze position along the saccade vector with an upsampling mechanism. This mechanism padded the number of data points along the saccade vector based on the fitted velocity profile. Microsaccades were discarded as biological implausible if the peak velocity of the upsampled saccade vector was three times higher (or more) than the peak velocity predicted for microsaccades of maximum amplitude (i.e., 1 dva) by the mean of the individual participant. In the final step, we inverted the coordinates of the upsampled saccade vector to replay the retinal consequences of the stimulus during a microsaccade (which shifts in the opposite direction of the eye movement). We applied the same data preprocessing in **Experiment 1** and **2**.

#### Exclusion of trials from analyses

Data obtained in the first session of the experiment were not considered in the main analysis, because the replay differed between this and later sessions of each participant. We also excluded trials in which a replayed microsaccade could render the stimulus visible or in which the participant generated at least one (additional) microsaccade. Trials in which a participant generated more than one microsaccade were likewise excluded from our analyses. Note that trials with accidental microsaccade generation in the fixation interval of **Experiment 1** were not excluded. We also disregarded trials with generated microsaccades larger than 1 dva and when the microsaccade failed to occur in the crucial time window of the stimulus (200–800 ms after stimulus onset.

### QUANTIFICATION AND STATISTICAL ANALYSIS

#### Motor control for microsaccades

##### Proportion of trials with microsaccade

We examined motor control for microsaccades by calculating the proportion of trials containing a microsaccade relative to the total number of trials. This was done separately for each eye movement type: intended and unintended microsaccades (**Exp. 1**), as well as spontaneous microsaccades (**Exp. 2**). Proportions were calculated individually per participant (irrespective of stimulus condition). To evaluate how well participants adjusted their eye movements to the different target distances (ranging from 0.2 to 1 dva), we categorized trials with intended microsaccades by target distances. While we did not preregister specific predictions, we expected a higher proportion of trials with microsaccade when participants were instructed to move their eyes (i.e., for intended microsaccades, **Exp. 1**), compared to when they were instructed to fixate (i.e., unintended microsaccades, **Exp 1**), or when they received no instruction.

To determine if observers generated more saccades when instructed to do so in **Experiment 1**, we calculated a paired two-sided t-test for the within-subject comparison of intended and all unintended microsaccades. To examine the effect of target distance, we conducted a one-way repeated-measures analysis of variance (rmANOVA) on the proportion of trials with intended microsaccades, categorized based on target distance. To compare the proportion of trials with saccades between experiments, we performed two two-sided independent samples t-tests, comparing spontaneous (**Exp. 2**) with unintended microsaccades (**Exp. 1**), as well as spontaneous (**Exp. 2**) and intended microsaccades (**Exp. 1**).

##### Saccade amplitudes

To further examine motor control of intended microsaccades, we calculated average amplitudes per participant for each of the five different target distances (0.2–1 dva). We did not pre-register specific hypotheses but would predict larger saccade amplitudes in trials with greater target distance.

To determine if more distant targets resulted in saccades with larger amplitudes, we conducted an rmANOVA with target amplitude (0.2–1.0 dva) as the within-subjects factor and saccade amplitude as the dependent variable to the intended microsaccades from **Experiment 1.** Individual means were calculated for each participant and each target amplitude level prior to analysis. We additionally fit a linear mixed effects model to the unaggregated amplitudes of the intended microsaccades. The model predicted saccade amplitude as a function of target distance using a restricted maximum likelihood (REML) method, with participants as random effects.

#### Visual sensitivity to intra-saccadic stimulation

##### Eye movement generation

We assessed observers’ visual stimulus sensitivity based on whether they provided a temporal estimate or pressed the space bar. A reported stimulus timing indicated stimulus perception, while pressing the space bar indicated no perception of the stimulus. Individual hit rates were calculated based on the number of trials with a stimulus and a temporal estimate. Similarly, individual false alarm rates were calculated based on temporal estimates in trials without a stimulus present. Since false alarm reports were rare, we chose not to calculate separate rates based on saccade generation and instead combined the rates for trials with and without eye movements. To assess whether stimulus visibility depended on the type of eye movement generated, we calculated different rates for trials with small intended and unintended saccades (**Exp. 1**), as well as spontaneous microsaccades (**Exp. 2**). To determine visual sensitivity for each participant and condition, individual hit and false alarm rates were z-transformed and subtracted. We predicted that visual sensitivity would depend on whether eye movements were generated, with sensitivity higher for trials with a microsaccade compared to trials without (i.e., generated microsaccades should yield higher sensitivity than non-saccadic trials). Conversely, we expected no difference in visual sensitivity between generated and replayed microsaccades. Lastly, we predicted that visual sensitivity would not differ based on the type of eye movement, meaning that intended, unintended, and spontaneous microsaccades would all produce similar sensitivity levels.

To determine whether the stimulus was invisible during stable fixation, we calculated average sensitivity indices for each eye movement type and compared their corresponding 95% confidence intervals (CI95%) to 0. Significant differences between eye movement types were assessed using paired two-sided t-tests for the within-subject comparison of intended and unintended eye movements (**Exp. 1**), and a two-sided independent samples t-test for the comparison between combined intended and unintended microsaccades (**Exp. 1**) and spontaneous microsaccades (**Exp. 2**). The effect of eye movement generation in **Experiment 1** was analyzed with a two-way repeated-measures ANOVA, using visual sensitivity indices as the dependent variable and stimulus condition (generated vs. replayed) and eye movement type (intended vs. unintended) as within-subject factors. To compare eye movements across experiments, we performed a mixed-measures ANOVA with stimulus condition (generated vs. replayed) as a within-subject factor and experiment (**Exp. 1** vs. **Exp. 2**) as a between-subject factor.

##### Eye movement kinematics

Since visual sensitivity is expected to depend on the degree of retinal stabilization of the stimulus, we calculated visual sensitivity as a function of retinal velocity in a second step. To determine retinal velocities, we subtracted the constant speed of the phase shift from the peak velocity of each microsaccade. We used directed speeds, with positive values for rightward and negative values for leftward oriented phase shifts and saccade directions. Retinal velocity of less than 30 dva/s were labelled ‘low’, while velocities above 30 dva/s were categorized as ‘high’. Hit and false alarm rates as well as visual sensitivities were calculated separately for generated and replayed eye movements and microsaccade types (intended and unintended microsaccades, **Exp. 1**; spontaneous microsaccades, **Exp. 2**), based on the resulting retinal velocity of the stimulus. We predicted that a higher retinal stability of the stimulus would lead to increased visual sensitivity. Consequently, microsaccades (irrespective of type) that lead to lower retinal velocities of the stimulus should result in higher sensitivity compared to trials in which the retinal velocity was high.

To assess if lower retinal stimulus velocities led to an increase in visual sensitivity in **Experiment 1**, we calculated a three-way rmANOVA with the within-subject factors retinal velocity (low vs. high velocity), stimulus condition (generated vs. replayed), and eye-movement type (intended vs. unintended). For the same analysis in **Experiment 2**, we conducted a two-way mixed-measures ANOVA with the within-subject factors retinal velocity (low vs. high) and stimulus condition (generated vs. replayed). Significant differences between factors were determined by calculating paired two-sided t-test for within-subject comparisons or two-sided independent samples t-test to compare between experiments whenever necessary.

One participant was excluded from this analysis due to the inability to compute a hit rate in a specific condition (i.e., unintended microsaccades associated with high retinal stimulus velocity).

#### Feeling of agency

To determine whether observers experience vFoA for microsaccades, we compared temporal estimates for generated and replayed eye movements. Temporal estimates were calculated based on the time point of highest stimulus visibility (t_visibility_). In active condition trials, this corresponded to the moment when a microsaccade reached its peak velocity, while in replay condition trials, it was the moment of highest aperture motion. This t_visibility_ was then subtract from the time of stimulus perception reported by the participant in the end of each trial *temporal estimate* = t*_report_* - t*_visitble_*) to obtain a positive estimate for trials in which participants overestimated the time point of stimulus appearance (i.e., t*_report_* - t*_visitble_*) and a negative estimate if stimulus timing was underestimated (i.e., t*_report_* - t*_visitble_*). We predicted that if participants experience a feeling of agency (FoA) for microsaccades, they would exhibit intentional binding and report the perceptual consequence of a microsaccade earlier in time compared to when the retinal consequence of the microsaccade was replayed. In other words, temporal reports should be significantly earlier in trials where a microsaccade made the stimulus visible, compared to trials where its visibility was caused by the replay of its retinal consequence. Importantly, we expected this effect only when the action was preceded by a corresponding intention. Thus, participants should report the retinal consequence of a microsaccade earlier in time (compared to its replay) only for intended microsaccades (**Exp. 1**), but not for unintended (**Exp. 1**) or spontaneous (**Exp. 2**) microsaccades.

To determine statistical significance, we used linear mixed-effects models (LMEs) to predict time intervals as a function of stimulus condition (active vs. replayed microsaccade), separately for each eye movement type (intended, unintended, spontaneous). The models included participants as random effects and allowed random intercepts and slopes:

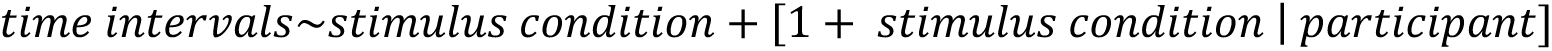

Additionally, we calculated Bayes Factors (BF_10_) to evaluate the relative evidence supporting or opposing our test hypothesis that there was a difference in time estimates between stimulus conditions (active vs. replay) for the different microsaccade types (intended, unintended, spontaneous). The analysis was conducted using a linear model with subjects as random effects. The null model assumed that any variation in time estimates was solely attributable to differences between subjects, independent of the stimulus condition.

Three participants were excluded from the analysis of unintended microsaccades because the model could not estimate the effect due to an insufficient number of temporal estimates.

To examine whether the amplitude of specifically intended eye movements influenced vFoA, we repeated the analysis described above for intended microsaccades generated at each target distance. While we did not pre-register specific predictions for this analysis, but expected vFoA would be more pronounced for larger saccade amplitudes.

## SUPPLEMENTARY MATERIAL

### Parameters of different types of generated and replayed microsaccades

**Fig. S1.**
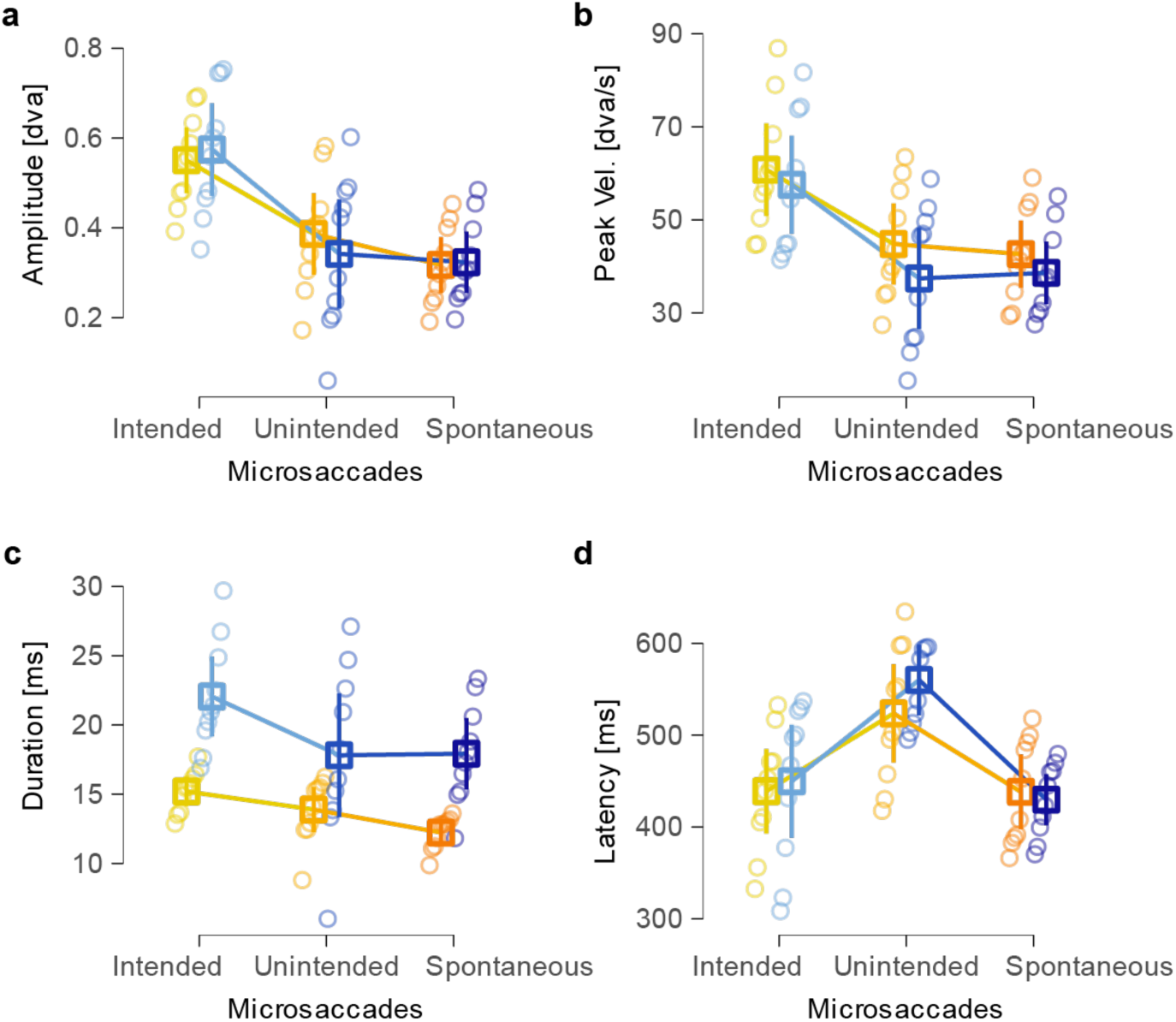
Parameters of unintended and spontaneous microsaccades are more similar for generated microsaccades, and intended and unintended microsaccades show greater similarity for replayed eye movements. **a** Comparison of amplitudes between intended, unintended (**Exp. 1**), and spontaneous microsaccades (**Exp. 2**) for generated (blues) and replayed (yellows) eye movements. **b** Comparison of peak velocities between eye movement types (same as in a). **c** Comparison of duration between eye movement types (same as in a). **d** Comparison of latencies between eye movement types (same as in a). In all plots, small circles indicate individual observers’ means, squares represent sample means. Error bars indicate 95% confidence intervals.

We compared amplitude, peak velocity, duration, and latency for intended, unintended (**Exp. 1**), and spontaneous microsaccades (**Exp. 2**). Starting with amplitude, we found the largest amplitudes for intended microsaccades (0.57±0.10 dva), whereas unintended microsaccades were significantly smaller (0.34±0.12 dva, *t* (9) = 3.33, *<u>p</u>* = 0.009) and comparable in size to spontaneous microsaccades from **Experiment 2** (0.32±0.07 dva, *t* (14.1) = 0.31, *p* > 0.250; **Fig. S1a**). For peak velocity, the results were similar: intended microsaccades yielded the highest peak velocities (57.54±10.55 dva/s), while the peak velocity of unintended microsaccades was significantly lower (37.41±10.89 dva/s; *t* (9) = 3.22, *p* = 0.01) but in the same range as spontaneous microsaccades (38.61±6.71 dva/s, *t* (15.0) = –0.21, *p* > 0.250; **Fig. S1b**). Microsaccade durations were, again, longest for intended eye movements (22.05±2.88 ms) with significantly slower unintended (17.81±4.47 ms, *t* (9) = 2.78, p = 0.021) and spontaneous eye movements from **Experiment 2** (17.93±2.57 ms, *t* (14.4) = –0.05, *p* > 0.250; **Fig. S1c**). Only saccade latencies deviated slightly from this pattern: they were longest for unintended movements from **Experiment 1** (560.17±38.56 ms)—significantly longer than both intended (449.84±61.57 ms, *t* (9) = –3.2, *p* = 0.011) and spontaneous ones (429.81±27.88 ms, *t* (16.4) = 6.2 < 0.001). Latencies between intended and spontaneous microsaccades did not differ significantly from each other (*t* (12.5) = 0.67 *p* > 0.250; **Fig. S1d**). These findings demonstrate that, overall, unintended and spontaneous microsaccades were comparable but different from intended eye movements.

For replayed microsaccades, a similar pattern emerged as for actively generated saccades: We found the longest amplitudes for intended eye movements (0.55±0.07 dva), with significantly smaller amplitudes for unintended (0.39±0.09 dva; *t* (9) = 4.41, *p* = 0.002) and spontaneous microsaccades (0.32±0.06 dva; *t* (17.5) = 5.49, *p* < 0.001; **Fig. S1a**). Our findings for peak velocities of replayed saccades mirrored those for amplitude, with significantly higher peak velocities for replayed intended (60.78 ± 9.94 dva/s) than unintended (44.81 ±8.73 dva/s; *t* (9) = 4.81, *p* < 0.001) and spontaneous eye movements (42.64 ±7.20 dva/s; *t* (16.4) = 3.34, *p* = 0.004; **Fig. S1b**). Durations of replayed microsaccades showed a slightly different pattern than actively generated saccades: While we found that intended had the longest durations (15.20± 1.06 ms), they were only insignificantly longer than those of unintended microsaccades (13.89±1.63 ms; *t* (9) = 1.62, *p* = 0.140). Spontaneous microsaccades were characterized by significantly shorter durations than intended microsaccades (12.23±0.84 ms; *t* (17.1) = 4.96, *p* < 0.001), but not unintended ones (*t* (13.5) = 2.04, *p* = 0.06; **Fig. S1c**). Latencies of replayed microsaccades, in turn, closely matched those of generated microsaccades with intended (438.94±46.06 ms; *t* (9) = –4.03, *p* < 0.001) and spontaneous microsaccades (438.29±40.74 ms; *t* (16.8) = 2.87, *p* < 0.011) significantly shorter than unintended ones (523.70±53.58 ms; **Fig. S1d**).

When comparing the parameters of intended, unintended, and spontaneous microsaccades between generated and replayed conditions, only three comparisons reached significance (all other *p*’s > 0.064). Replayed intended and spontaneous microsaccades had significantly shorter durations than their generated counterparts (intended: *t* (9) = –5.93, *p* < 0.001; spontaneous: *t* (9) = –5.71, *p* < 0.001). Additionally, spontaneous microsaccades exhibited a slightly, yet significantly, lower peak velocity when replayed (*t* (9) = 3.62, *p* < 0.006). We therefore conclude that the replayed microsaccades closely matched the generated eye movements, allowing for a valid comparison of temporal differences between stimulus conditions to assess effect binding and, hence, the role of agency for visual consequences of eye movements.

## References

Blakemore, S.-J., Wolpert, D., & Frith, C. (2000). Why can’t you tickle yourself?: NeuroReport, 11(11), R11–R16. 10.1097/00001756-200008030-00002

Brainard, D. H. (1997). The Psychophysics Toolbox. Spatial Vision, 10(4), 433–436. 10.1163/156856897X00357

Bridgeman, B. (1995). A review of the role of efference copy in sensory and oculomotor control systems. Annals of Biomedical Engineering, 23(4), 409–422. 10.1007/BF02584441

Brown, P., & Rothwell, J. C. E. (1997). Illusions of time. Abstracts, 27th Annual Meeting, 23, 1119.

Castet, E., & Masson, G. S. (2000). Motion perception during saccadic eye movements. Nature Neuroscience, 3(2), 177–183. 10.1038/72124

Clark, A., Kiverstein, J., & Vierkant, T. (Eds.). (2013). Decomposing the will. Oxford University Press.

Collewijn, H., Erkelens, C. J., & Steinman, R. M. (1988). Binocular co-ordination of human horizontal saccadic eye movements. The Journal of Physiology, 404(1), 157–182. 10.1113/jphysiol.1988.sp017284

Cornelissen, F. W., Peters, E. M., & Palmer, J. (2002). The Eyelink Toolbox: Eye tracking with MATLAB and the Psychophysics Toolbox. Behavior Research Methods, Instruments, & Computers, 34(4), 613–617. 10.3758/BF03195489

Deubel, H., & Elsner, T. (1986). Threshold perception and saccadic eye movements. Biological Cybernetics, 54(6), 351–358. 10.1007/BF00355540

Deubel, H., Elsner, T., & Hauske, G. (1987). Saccadic eye movements and the detection of fast-moving gratings. Biological Cybernetics, 57(1–2), 37–45. 10.1007/BF00318714

Engbert, R., & Kliegl, R. (2003). Microsaccades uncover the orientation of covert attention. Vision Research, 43(9), 1035–1045. 10.1016/S0042-6989(03)00084-1

Engbert, R., & Mergenthaler, K. (2006). Microsaccades are triggered by low retinal image slip. Proceedings of the National Academy of Sciences, 103(18), 7192–7197. 10.1073/pnas.0509557103

Findlay, J. M. (1974). Direction perception and human fixation eye movements. Vision Research, 14(8), 703–711. 10.1016/0042-6989(74)90067-4

Gallagher, S. (2000). Philosophical conceptions of the self: Implications for cognitive science. Trends in Cognitive Sciences, 4(1), 14–21. 10.1016/S1364-6613(99)01417-5

Hafed, Z. M. (2011). Mechanisms for generating and compensating for the smallest possible saccades. European Journal of Neuroscience, 33(11), 2101–2113. 10.1111/j.1460-9568.2011.07694.x

Hafed, Z. M., & Goffart, L. (2020). Gaze direction as equilibrium: More evidence from spatial and temporal aspects of small-saccade triggering in the rhesus macaque monkey. Journal of Neurophysiology, 123(1), 308–322. 10.1152/jn.00588.2019

Hafed, Z. M., Goffart, L., & Krauzlis, R. J. (2009). A Neural Mechanism for Microsaccade Generation in the Primate Superior Colliculus. Science, 323(5916), 940–943. 10.1126/science.1166112

Haggard, P. (2017). Sense of agency in the human brain. Nature Reviews Neuroscience, 18(4), 196–207. 10.1038/nrn.2017.14

Haggard, P., & Clark, S. (2003). Intentional action: Conscious experience and neural prediction. Consciousness and Cognition, 12(4), 695–707. 10.1016/S1053-8100(03)00052-7

Hogendoorn, H. (2016). Voluntary Saccadic Eye Movements Ride the Attentional Rhythm. Journal of Cognitive Neuroscience, 28(10), 1625–1635. 10.1162/jocn_a_00986

Hunt, A. R., & Cavanagh, P. (2009). Looking ahead: The perceived direction of gaze shifts before the eyes move. Journal of Vision, 9(9), 1–1. 10.1167/9.9.1

Imaizumi, S., & Tanno, Y. (2019). Intentional binding coincides with explicit sense of agency. Consciousness and Cognition, 67, 1–15. 10.1016/j.concog.2018.11.005

Kelly, D. H. (1990). Moving gratings and microsaccades. Journal of the Optical Society of America A, 7(12), 2237. 10.1364/JOSAA.7.002237

Kennard, C., Mannan, S. K., Nachev, P., Parton, A., Mort, D. J., Rees, G., Hodgson, T. L., & Husain, M. (2005). Cognitive Processes in Saccade Generation. Annals of the New York Academy of Sciences, 1039(1), 176–183. 10.1196/annals.1325.017

Klanke, J.-N., Ohl, S., & Rolfs, M. (2024). Sensorimotor awareness requires intention: Evidence from minuscule eye movements. Neuroscience. 10.1101/2024.07.02.601661

Kleiner, M., Brainard, D. H., Pelli, D., Ingling, A., Murray, R., & Broussard, C. (2007). What’s new in psychtoolbox-3. Perception, 36(14), 1–16.

Ko, H., Poletti, M., & Rucci, M. (2010). Microsaccades precisely relocate gaze in a high visual acuity task. Nature Neuroscience, 13(12), 1549–1553. 10.1038/nn.2663

Legaspi, R., & Toyoizumi, T. (2019). A Bayesian psychophysics model of sense of agency. Nature Communications, 10(1), 4250. 10.1038/s41467-019-12170-0

Martinez-Conde, S., Macknik, S. L., & Hubel, D. H. (2004). The role of fixational eye movements in visual perception. Nature Reviews Neuroscience, 5(3), 229–240. 10.1038/nrn1348

Martinez-Conde, S., Otero-Millan, J., & Macknik, S. L. (2013). The impact of microsaccades on vision: Towards a unified theory of saccadic function. Nature Reviews Neuroscience, 14(2), 83–96. 10.1038/nrn3405

Moore, J. W., & Obhi, S. S. (2012). Intentional binding and the sense of agency: A review. Consciousness and Cognition, 21(1), 546–561. 10.1016/j.concog.2011.12.002

Morrone, M. C., Ross, J., & Burr, D. (2005). Saccadic eye movements cause compression of time as well as space. Nature Neuroscience, 8(7), 950–954. 10.1038/nn1488

Otero-Millan, J., Macknik, S. L., & Martinez-Conde, S. (2014). Fixational eye movements and binocular vision. Frontiers in Integrative Neuroscience, 8. 10.3389/fnint.2014.00052

Otero-Millan, J., Troncoso, X. G., Macknik, S. L., Serrano-Pedraza, I., & Martinez-Conde, S. (2008). Saccades and microsaccades during visual fixation, exploration, and search: Foundations for a common saccadic generator. Journal of Vision, 8(14), 21–21. 10.1167/8.14.21

Pashler, H. (1994). Dual-task interference in simple tasks: Data and theory. Psychological Bulletin, 116(2), 220–244. 10.1037/0033-2909.116.2.220

Pelli, D. G. (1997). The VideoToolbox software for visual psychophysics: Transforming numbers into movies. Spatial Vision, 10(4), 437–442. 10.1163/156856897X00366

Poletti, M. (2023). An eye for detail: Eye movements and attention at the foveal scale. Vision Research, 211, 108277. 10.1016/j.visres.2023.108277

Poletti, M., Intoy, J., & Rucci, M. (2020). Accuracy and precision of small saccades. Scientific Reports, 10(1), 16097. 10.1038/s41598-020-72432-6

Rolfs, M. (2009). Microsaccades: Small steps on a long way. Vision Research, 49(20), 2415– 2441. 10.1016/j.visres.2009.08.010

Rolfs, M., Kliegl, R., & Engbert, R. (2008). Toward a model of microsaccade generation: The case of microsaccadic inhibition. Journal of Vision, 8(11), 5–5. 10.1167/8.11.5

Rucci, M., & Poletti, M. (2015). Control and Functions of Fixational Eye Movements. Annual Review of Vision Science, 1(1), 499–518. 10.1146/annurev-vision-082114-035742

Ruess, M., Thomaschke, R., & Kiesel, A. (2020a). Acting and reacting: Is intentional binding due to sense of agency or to temporal expectancy? Journal of Experimental Psychology: Human Perception and Performance, 46(1), 1–9. 10.1037/xhp0000700

Ruess, M., Thomaschke, R., & Kiesel, A. (2020b). Intentional Binding for Unintended Effects. Timing & Time Perception, 8(3–4), 341–349. 10.1163/22134468-bja10005

Schwarz, K. A., & Weller, L. (2023). Distracted to a fault: Attention, actions, and time perception. *Attention, Perception*, & Psychophysics, 85(2), 301–314. 10.3758/s13414-022-02632-x

Schweitzer, R., & Rolfs, M. (2022). Definition, Modeling, and Detection of Saccades in the Face of Post-saccadic Oscillations. In S. Stuart (Ed.), Eye Tracking (Vol. 183, pp. 69– 95). Springer US. 10.1007/978-1-0716-2391-6_5

Skavenski, A. A., Haddad, G., & Steinman, R. M. (1972). The extraretinal signal for the visual perception of direction. Perception & Psychophysics, 11(4), 287–290. 10.3758/BF03210380

Steinbach, M. J. (1987). Proprioceptive knowledge of eye position. Vision Research, 27(10), 1737–1744. 10.1016/0042-6989(87)90103-9

Suzuki, K., Lush, P., Seth, A. K., & Roseboom, W. (2019). Intentional Binding Without Intentional Action. Psychological Science, 30(6), 842–853. 10.1177/0956797619842191

Synofzik, M., Vosgerau, G., & Newen, A. (2008). Beyond the comparator model: A multifactorial two-step account of agency. Consciousness and Cognition, 17(1), 219–239. 10.1016/j.concog.2007.03.010

Willeke, K. F., Tian, X., Buonocore, A., Bellet, J., Ramirez-Cardenas, A., & Hafed, Z. M. (2019). Memory-guided microsaccades. Nature Communications, 10(1), 3710. 10.1038/s41467-019-11711-x

Wolfe, J. M., Alvarez, G. A., Rosenholtz, R., Kuzmova, Y. I., & Sherman, A. M. (2011). Visual search for arbitrary objects in real scenes. Attention, Perception, & Psychophysics, 73(6), 1650–1671. 10.3758/s13414-011-0153-3

World Medical Association Declaration of Helsinki: Ethical Principles for Medical Research Involving Human Subjects. (2013). JAMA, 310(20), 2191. 10.1001/jama.2013.281053

Yarrow, K., Haggard, P., Heal, R., Brown, P., & Rothwell, J. C. (2001). Illusory perceptions of space and time preserve cross-saccadic perceptual continuity. Nature, 414(6861), 302–305. 10.1038/35104551

Yarrow, K., Whiteley, L., Haggard, P., & Rothwell, J. C. (2006). Biases in the perceived timing of perisaccadic perceptual and motor events. Perception & Psychophysics, 68(7), 1217–1226. 10.3758/BF03193722

Zuber, B. L., Stark, L., & Cook, G. (1965). Microsaccades and the Velocity-Amplitude Relationship for Saccadic Eye Movements. Science, 150(3702), 1459–1460. 10.1126/science.150.3702.1459

